# Unlocking the Role of Endothelial MPL Receptor and JAK2V617F Mutation: Insights into Cardiovascular Dysfunction in MPNs and CHIP

**DOI:** 10.1101/2023.07.12.548716

**Authors:** Haotian Zhang, Nicholas Kafeiti, Sandy Lee, Kyla Masarik, Haoyi Zheng, Huichun Zhan

## Abstract

Patients with JAK2V617F-positive myeloproliferative neoplasms (MPNs) and clonal hematopoiesis of indeterminate potential (CHIP) are at a significantly higher risk of cardiovascular diseases (CVDs). Endothelial cells (ECs) carrying the JAK2V617F mutation can be detected in many MPN patients. Here, we investigated the impact of endothelial JAK2V617F mutation on CVD development using both transgenic murine models and human induced pluripotent stem cell lines. Our findings revealed that JAK2V617F mutant ECs promote CVDs by impairing endothelial function and undergoing endothelial-to-mesenchymal transition (EndMT). Importantly, we found that inhibiting the endothelial thrombopoietin receptor MPL suppressed JAK2V617F-induced EndMT and prevented cardiovascular dysfunction caused by mutant ECs. These findings propose that targeting the endothelial MPL receptor could be a promising therapeutic approach to manage CVD complications in patients with JAK2V617F-positive MPNs and CHIP. Further investigations into the impact of other CHIP-associated mutations on endothelial dysfunction are needed to improve risk stratification for individuals with CHIP.

## Introduction

The acquired kinase mutation JAK2V617F is one of the common mutations associated with clonal hematopoiesis of indeterminate potential (CHIP), and can be detected in ∼0.2% individuals of the general population^1,2^. Individuals with CHIP have a 2-4 fold increased risk of cardiovascular diseases (CVDs) with worsened clinical outcomes, which rivals or even exceeds those traditional CVD risk factors such as diabetes, hypertension, hyperlipidemia, and smoking^3-7^. In particular, individuals with JAK2V617F mutant CHIP have a 12-fold higher risk of coronary heart disease and ischemic stroke compared to individuals without any CHIP-associated mutation^5,7,8^. Even a small amount of JAK2V617F mutation (with allele frequency 0.06-1.7%) is associated with more than a 5-fold increased risk of CVDs compared to individuals without the mutation^8^. JAK2V617F also plays a central role in most patients with myeloproliferative neoplasms (MPNs), a group of clonal stem cell disorders characterized by hematopoietic stem/progenitor cell (HSPC) expansion and overproduction of mature, often dysfunctional blood cells. Up to 40-50% of patients with MPNs experience arterial or venous thrombosis during the course of their disease, with CVDs being the leading cause of morbidity and mortality in these patients, a markedly increased incidence compared to age-matched controls^9-11^. Given that up to 50% of MPN patients develop thrombosis and CVDs are the leading cause of morbidity and mortality in these patients, and the presence of MPN-associated mutation in seemingly healthy individuals is linked to an excess of cardiovascular risks, JAK2V617F-positive MPNs provide an excellent model system to investigate the relationship between hematopoietic mutations and CVDs.

Vascular endothelial cells (ECs) play a critical role in regulating cardiovascular function, and their dysfunction can lead to micro- and macro-vascular complications in CVDs. While the common origin of blood and endothelium beyond embryonic development is still under investigation, hematopoietic mutations have been detected in ECs in many hematologic malignancies, including MPNs, lymphoma, and myeloma^12-19^. Specifically, the JAK2V617F mutation has been identified in microvascular ECs isolated from the liver^12^, spleen^13^ (using laser microdissection), and bone marrow^16^ (using flow cytometry) of MPNs patients. Moreover, this mutation has been found in EC progenitors derived from the hematopoietic lineage and, in some reports based on *in vitro* assays, in true endothelial colony-forming cells from patients with MPNs^13,17,18,20,21^.

In our previous study, we demonstrated that a murine model of JAK2V617F-positive MPN, in which the mutation is expressed in both blood cells and vascular ECs, develops spontaneous heart failure with a thrombosis and vasculopathy phenotype^22^. Unlike other murine models of CHIP mutations (e.g., TET2, JAK2V617F), in which CVDs only develop in the presence of additional risk factors (e.g., high-fat diet^7,23^, surgical constriction of coronary artery or aorta^24,25^, or hyperlipidemia^26^), the JAK2V617F-positive mice with both mutant blood cells and mutant vascular ECs develop spontaneous CVD even when fed a regular chow diet, suggesting that mutant ECs can accelerate cardiovascular dysfunction. In this study, we aimed to investigate the mechanisms through which the JAK2V617F mutation induces endothelial dysfunction to promote CVDs using both transgenic murine models and induced pluripotent stem cell (iPS) lines derived from an MPN patient.

## Results

### Both mutant ECs and mutant blood cells are required to drive spontaneous cardiovascular dysfunction in a JAK2V617F-positive MPN murine model

Previously, we crossed mice that bear a Cre-inducible human JAK2V617F gene (FF1)^27^ with Tie2-Cre mice^28^ to express JAK2V617F specifically in all hematopoietic cells and ECs (Tie2-cre^+/-^FF1^+/-^, or Tie2^+^FF1^+^). Tie2^+^FF1^+^ mice developed MPN and spontaneous thrombosis, vasculopathy, and cardiomyopathy phenotype with an increased risk of sudden death^22^. We also demonstrated that the presence of JAK2V617F-mutant blood cells alone was insufficient to generate the spontaneous CVD phenotype; mutant ECs were required for the development of this phenotype^22^. To further investigate the impact of JAK2V617F mutant ECs on CVD development, we crossed FF1 mice with VEcadherin-creERT2 mice^29,30^ (which express the Cre recombinase specifically in ECs upon tamoxifen induction) to generate VEcad-cre^+/-^FF1^+/-^ mice (or CDH5^+^FF1^+^). The efficiency of endothelial recombination and the absence of hematopoietic recombination were previously validated^30,31^. At approximately 6-8 weeks of age, CDH5^+^FF1^+^ mice were treated with 10mg tamoxifen^29,30^ to induce the expression of human JAK2V617F specifically in vascular ECs but not in blood cells (Figure 1A). Twelve weeks after tamoxifen induction, CDH5^+^FF1^+^ mice exhibited mild thrombocytosis, but their hemoglobin and white blood cell counts remained normal (Figure 1B), resembling the blood profile of patients with mild essential thrombocythemia. No splenomegaly was observed in CDH5^+^FF1^+^ mice compared to age-matched control mice (Figure 1C). These results are consistent with a previous study that used the same VEcadherin-creERT2 mouse but employed a different JAK2V617F transgenic mouse^31^. Serial transthoracic echocardiography demonstrated no significant alterations in left ventricular (LV) ejection fraction or volume between CDH5^+^FF1^+^ mice and age-matched control mice (Figure 1D). Additionally, there were no significant differences in heart weight between CDH5^+^FF1^+^ mice and control mice (Figure 1E).

**Figure 1.**
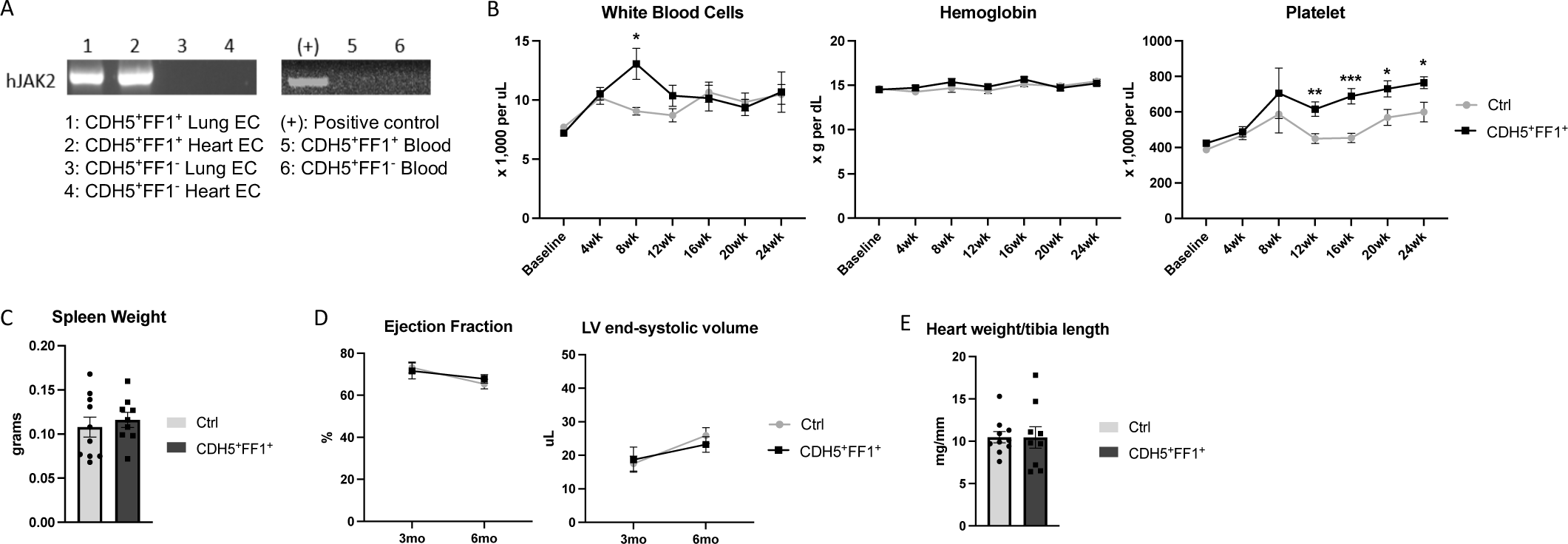
Effects of JAK2V617F mutation in ECs on both hematologic and cardiovascular phenotypes. (**A**) Expression of human JAK2 gene in isolated lung and heart ECs (left) and peripheral blood cells (right) from CDH5^+^FF1^+^ mice and control mice 6 weeks after tamoxifen induction. (**B**) Peripheral blood counts in CDH5^+^FF1^+^ and control mice after tamoxifen induction (n=6-22 mice in each group). (**C**) Spleen weight of CDH5^+^FF1^+^ and control mice 6-7 months after tamoxifen induction (n=9-10 mice in each group). (**D**) Serial measurements of cardiac ejection fraction and LV end-systolic volume in CDH5^+^FF1^+^ mice and control mice (n=6 mice in each group). (**E**) Heart weight adjusted by tibia length in CDH5^+^FF1^+^ mice and control mice 6 months after tamoxifen induction (n=9-10 mice in each group).

Taken together with our previous findings^22^, these results indicate that both mutant ECs and mutant blood cells are required for the development of a spontaneous CVD phenotype in the JAK2V617F-positive murine model of MPNs.

### The JAK2V617F mutant ECs promote CVDs when challenged with a high-fat diet treatment

To test whether JAK2V617F mutant ECs can promote the development of CVDs when challenged with additional stressors, we treated both CDH5^+^FF1^+^ mice and VEcadherin-creERT2 control mice (CDH5^+^FF1^-^) with a high-fat/high-cholesterol diet (TD.88137, Harlan-Tekla®). After a 6-week diet treatment, CDH5^+^FF1^+^ mice exhibited a phenotype of dilated cardiomyopathy, characterized by significant decreases in LV ejection fraction and increases in LV end-diastolic and end-systolic volumes compared to control mice. These phenotypic differences continued to worse following a 12-week diet treatment (Figure 2A).

**Figure 2.**
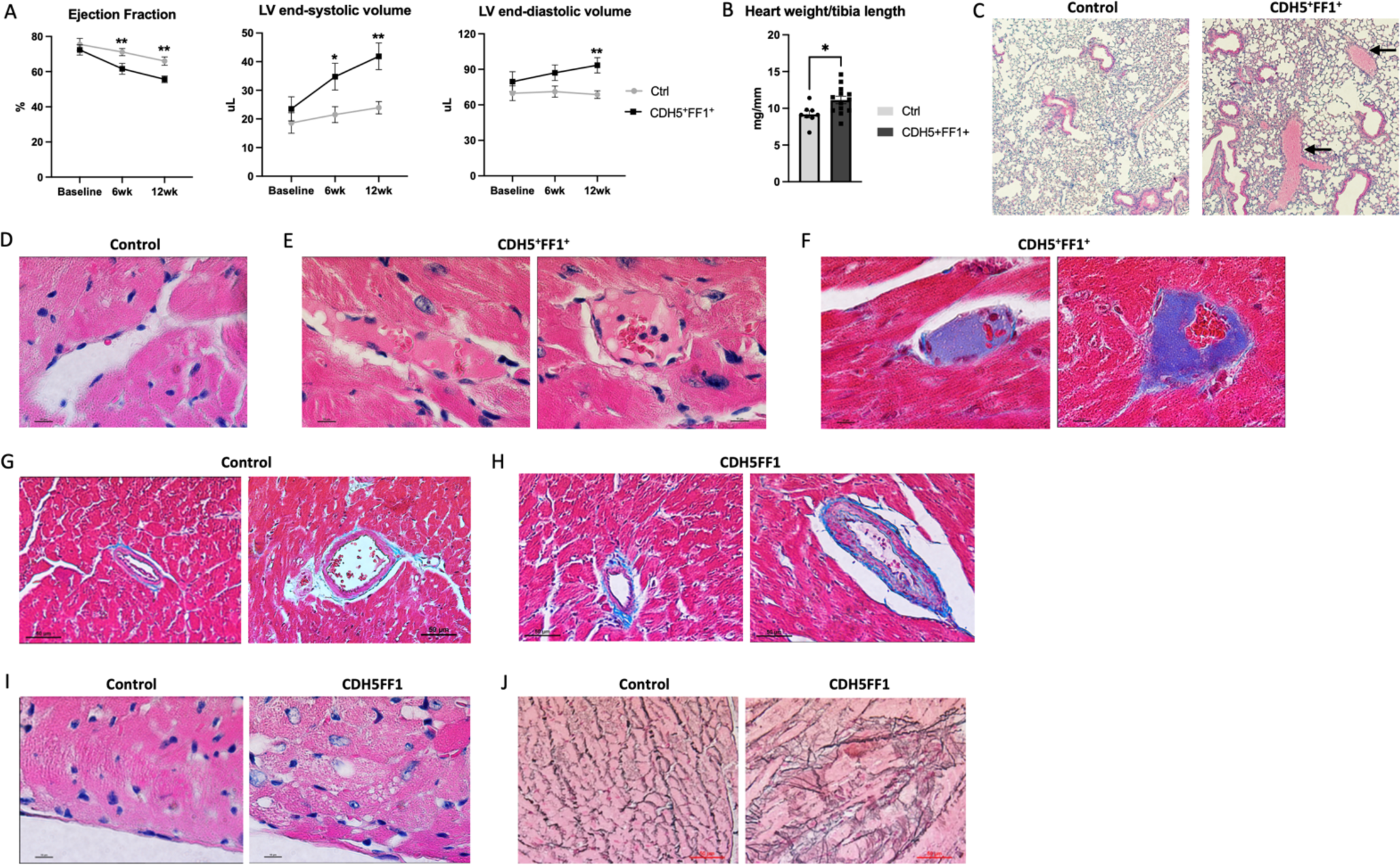
The JAK2V617F mutant ECs promote CVDs when challenged with a high-fat diet treatment. (**A**) LV ejection fraction and LV volumes in CDH5^+^FF1^+^ and control mice after high-fat diet challenge (n=8-13 mice in each group). (**B**) Heart weight adjusted by tibia length in CDH5^+^FF1^+^ and control mice after 12-week high-fat diet (HFD) treatment (n=8-13 mice in each group). (**C**) H&E staining of lung sections from control and CDH5^+^FF1^+^ mice after 12-week HFD treatment. Arrows indicate thrombi in segment pulmonary arteries of CDH5^+^FF1^+^ mice (magnification 10x). (**D**) H&E staining of a coronary capillary in control mice (magnification 100x). (**E-F**) H&E staining (E) and Masson’s trichrome staining (F) showed fibrin clots in scattered coronary capillaries in the CDH5^+^FF1^+^ mice (magnification 100x). (**G-H**) Masson’s trichrome staining of coronary arterioles in control mice (G) and CDH5^+^FF1^+^ mice (H). There was intimal thickening, smooth muscle layer thickening, and perivascular fibrosis of coronary arterioles in the CDH5^+^FF1^+^ mice compared to control mice (magnification 40x). (**I**) H&E-stained cardiac sections (taken from similar locations of the heart) reveal signs of cardiomyocyte damage in CDH5^+^FF1^+^ mice, as suggested by the distortion to the normal architecture, clearing of the cytoplasm, and irregular cell borders. (**J**) Reticulin staining of cardiac sections (taken from similar locations of the heart) from control and CDH5^+^FF1^+^ mice after 12-wk HFD treatment (magnification 40x). Note: Images are representative examples. N = 4 mice in each group were examined.

Pathological evaluation confirmed the diagnosis of dilated cardiomyopathy in CDH5^+^FF1^+^ mice, as evidenced by a significantly increased heart weight-to-tibia length ratio compared to controls (Figure 2B). Thromboses were observed in segment pulmonary arteries and scattered coronary capillaries, whereas age-matched control mice showed no evidence of spontaneous thrombosis in the heart or lungs (Figure 2C-F). Further examination of the coronary vasculature revealed intimal thickening, smooth muscle cell thickening, and perivascular fibrosis of scattered coronary arterioles in CDH5^+^FF1^+^ mice compared to control mice (Figure 2G-H). Additionally, there were signs of cardiomyocyte damage (e.g., distortion of the normal architecture, cytoplasm clearing, and irregular cell borders) (Figure 2I) and increased reticulin fibers (Figure 2J) in the heart of CDH5^+^FF1^+^ mice following high-fat diet challenge, indicating cardiac remodeling resulting from microvascular thrombosis and vasculopathy/vascular remodeling. Notably, no atherosclerotic lesions were observed.

Overall, these findings indicate that a high-fat diet challenge promotes cardiovascular dysfunction in the presence of mutant ECs alone, highlighting the critical role of endothelial dysfunction in the development of cardiovascular complications in JAK2V617F-positive MPN murine models.

### The JAK2V617F mutation promotes endothelial-to-mesenchymal transition and endothelial dysfunction in transgenic murine models

We then investigated the impact of high-fat diet treatment on the function of JAK2V617F mutant ECs. First, under regular diet conditions, JAK2V617F mutant cardiac ECs (CD45^-^CD31^+^) exhibited moderately increased angiogenesis *in vitro* compared to wild-type cardiac ECs (Figure 3A). However, when exposed to a high-fat diet, JAK2V617F mutant cardiac ECs demonstrated significantly reduced angiogenesis compared to wild-type ECs (Figure 3B). We then examined the expression levels of vascular endothelial-cadherin (VE-cadherin)^32^ and VEGF-R2^33^, two key endothelial cell proteins involved in angiogenesis and vascular integrity. Quantitative flow cytometry analysis revealed no difference in VE-cadherin or VEGF-R2 expression between wild-type cardiac ECs and JAK2V617F mutant cardiac ECs from mice on a regular diet (Figures 3C-D). However, their expression was significantly decreased in JAK2V617F mutant ECs compared to wild-type cardiac ECs from mice following the high-fat diet treatment (Figures 3E-F).

**Figure 3.**
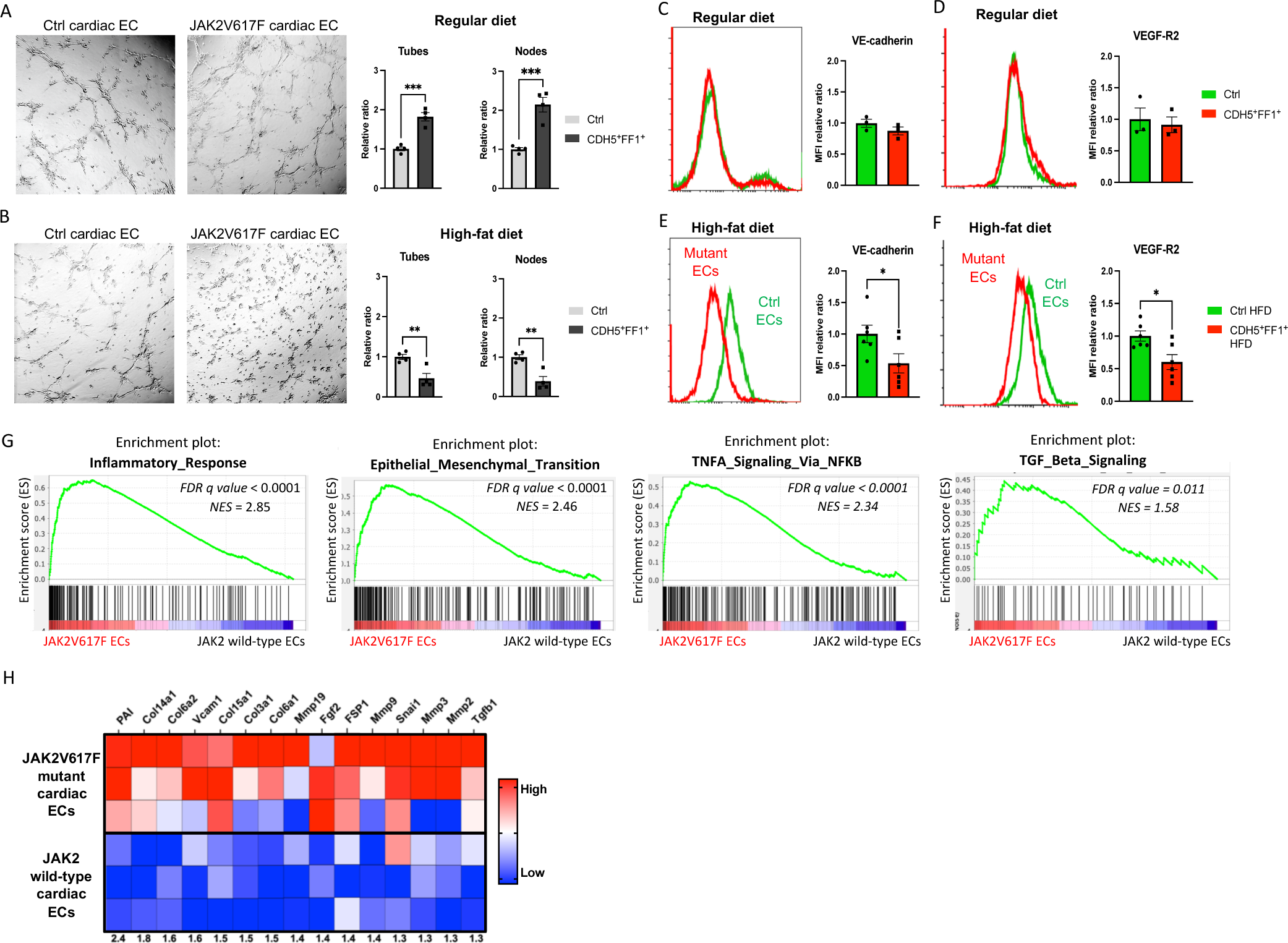
JAK2V617F mutant ECs display endothelial dysfunction with increased markers of endothelia-to-mesenchymal transition. (**A-B**) Tube formation assays showing increased tube formation in JAK2V617F mutant cardiac ECs compared to wild-type cardiac ECs from control mice on a regular chow diet (A) and decreased tube formation in JAK2V617F mutant cardiac ECs compared to wild-type cardiac ECs after high-fat diet treatment (B). Quantification of tube formation was performed on images taken at 4x magnification by counting the number of tubes in four non-overlapping fields. Results are expressed as the mean ± standard error of the mean (n = 4). Data are from one of three independent experiments that gave similar results. (**C-F**) Expression levels of VE-cadherin and VEGF-R2 expression in JAK2V617F mutant and wild-type cardiac ECs during regular diet (C-D) and high-fat diet (E-F) treatment. n = 3 mice in the regular diet groups (C-D) and n = 6 mice in the high-fat diet group (E-F). Similar results were obtained from two independent experiments. (**G**) Gene set enrichment analysis (GSEA) of differentially regulated genes in wild-type cardiac ECs (from control mice with normal cardiac function) and JAK2V617F mutant cardiac ECs (from Tie2^+^FF1^+^ mice with CVD). Both inflammatory response genes and EndMT genes are significantly enriched in JAK2V617F mutant cardiac ECs. (**H**) Heatmaps showing the expression of EndMT genes in wild-type and JAK2V617F mutant cardiac ECs.

In a previous study, we conducted whole-transcriptome RNA sequencing and identified gene expression changes associated with the JAK2V617F mutation in murine cardiac ECs^22^. Notably, gene expression signatures related to inflammatory response, epithelial-mesenchymal transition, tumor necrosis factor-α (TNFα) signaling, and transforming growth factor-? (TGF?) signaling were the most significantly upregulated in JAK2V617F mutant cardiac ECs from Tie2^+^FF1^+^ mice (which developed spontaneous CVD) compared to wild-type cardiac ECs from control mice (Figure 3G). Additionally, genes associated with endothelial-to-mesenchymal transition (EndMT) were significantly upregulated in JAK2V617F mutant cardiac ECs (Figure 3H), including collagen genes such as COL3A1 and COL6A1/2, matrix metalloproteinase genes such as MMP2 and MMP3, cell adhesion genes such as VCAM1, EndMT-associated signaling genes such as TGF?1 and FGF2, as well as several EndMT-associated markers such as Plasminogen activator inhibitor 1 (PAI) and Fibroblast-specific protein 1 (FSP1)^34,35^.

EndMT is a key process involved in normal heart development and can be induced postnatally, contributing to endothelial dysfunction and various cardiovascular diseases^34-36^. During EndMT, endothelial cells lose their EC markers and functions (e.g., angiogenesis) and acquire mesenchymal cell characteristics. Inflammation is a well-known inducer of EndMT, and the TGF? and TNFα signaling pathways play significant roles in inflammation-induced EndMT^34-36^. We have observed consistent findings across different experimental models, including the presence of coronary vasculopathy and cardiac fibrosis in both Tie2^+^FF1^+^ mice (with spontaneous CVD on a regular diet)^22^ and CDH5^+^FF1^+^ mice (with high-fat diet-induced CVD) (Figure 2), impaired EC function and loss of EC markers in JAK2V617F mutant cardiac ECs exposed to a high-fat diet (Figure 3A-F), and the significant enrichment of genes associated with EndMT in JAK2V617F mutant cardiac ECs from mice with CVD (Figure 3G-H). These findings provide compelling evidence for the involvement of EndMT in JAK2V617F-associated cardiovascular complications.

### The JAK2V617F mutation induces endothelial-to-mesenchymal transition in induced pluripotent stem (iPS) cell lines derived from an MPN patient

The study of human disease is often limited by the availability of relevant cells and tissues for study. Primary HSPCs and ECs derived from patients often consist of a heterogeneous mixture of normal and diseased cells, presenting difficulties in functional analysis due to genetic variability. To overcome these challenges, we utilized a pair of isogenic iPS cell lines derived from the same MPN patient, one carrying the JAK2V617F mutation (PVB1.4) and the other being JAK2 wild-type (PVB1.11)^37^. These iPS cell lines provided a unique opportunity to investigate the impact of the JAK2V617F mutation on human endothelial function under a controlled genetic background.

We successfully differentiated both JAK2 wild-type (PVB1.11) and JAK2V617F mutant (PVB1.4) iPS cell lines into ECs using a modified iPS differentiation protocol^38,39^ (Figure 4A). After a 6-day differentiation process, we observed similar expression of EC markers (e.g., CD31, CD34) in both PVB1.11 and PVB1.4 cell lines (Figure 4B). These findings suggest that the presence of the JAK2V617F mutation did not hinder the differentiation of iPS cells into ECs.

**Figure 4.**
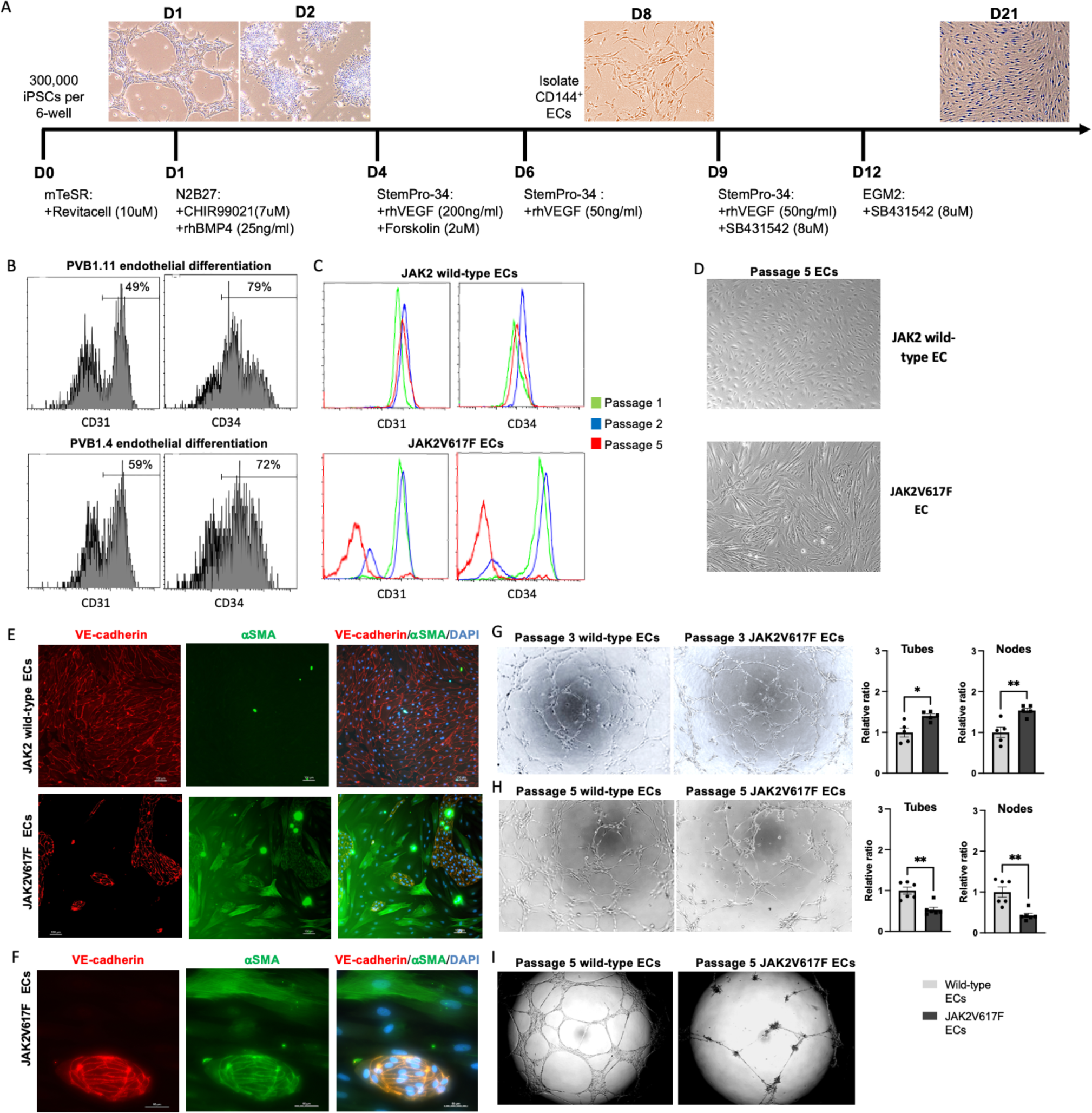
The JAK2V617F mutation induces endothelial-to-mesenchymal transition in iPS cell lines derived from an MPN patient. (**A**) Schematic presentation of the EC differentiation strategy for human iPS cells. (**B**) Flow cytometry analysis showing the expression of endothelial markers CD31 and CD34 on cells differentiated from PVB1.11 (JAK2 wild-type; top) and PVB1.4 (JAK2V617F mutant; bottom) iPS lines after 6 days of EC differentiation. (**C**) Flow cytometric characterization of human iPS-derived ECs during continuous *in vitro* culture. The expression of CD31 and CD34 was analyzed on passages 1, 2, and 5 ECs derived from PVB1.11 (top) and PVB1.4 (bottom) iPS lines. Similar results were obtained from two independent experiments performed by two different researchers. (**D**) Representative bright field images of passage 5 wild-type (top) and JAK2V617F mutant (bottom) ECs, showing their morphological differences (magnification: 10x). (**E**) Immunofluorescence staining for the endothelial marker VE-cadherin (red) and smooth muscle cell marker αSMA (green) in passage 4 ECs derived from PVB1.11 (top) and PVB1.4 (bottom) iPS lines. (**F**) Co-expression of both VE-cadherin and αSMA in some passage 4 ECs derived from PVB1.4 iPS line. (**G-H**) Tube formation assays of passage 3 (G) and passage 5 (H) JAK2V617F mutant ECs compared to wild-type ECs after a 2-hour incubation in Matrigel matrix (magnification: 10x). (**I**) Stability of tubes formed by passage 5 JAK2V617F ECs compared to wild-type ECs after a 24-hour incubation in the Matrigel matrix (magnification: 4x).

We then isolated CD144+ ECs derived from the iPS cell lines and conducted continuous *in vitro* culture. JAK2 wild-type ECs derived from PVB1.11 maintained the expression of EC markers throughout serial passages. In contrast, JAK2V617F mutant ECs derived from PVB1.4 exhibited a gradual loss of CD31 and CD34 expression (Figures 4C). Moreover, these mutant EC underwent a phenotypic transition toward a fibroblastoid morphology, characterized by an elongated shape (Figure 4D), indicating a transition towards a mesenchymal phenotype. Immunofluorescence staining confirmed a significant decrease in the expression of the endothelial marker VE-cadherin in PVB1.4-derived ECs compared to PVB1.11-derived ECs. Additionally, we observed the emergence of smooth muscle actin (αSMA)-positive cells within the PVB1.4-derived EC culture (Figure 4E). Notably, some PVB1.4-derived ECs displayed co-expression of both VE-cadherin and αSMA, suggesting a partial acquisition of smooth muscle cell characteristics (Figure 4F).

Functional analysis further revealed that, while earlier passage PVB1.4 ECs displayed increased angiogenesis compared to PVB1.11 ECs (Figure 4G), passage 5 PVB1.4 ECs exhibited significantly decreased angiogenesis (Figure 4H) with less stable tube formation (Figure 4I).

Collectively, these findings strongly support the notion that JAK2V617F mutant human ECs derived from an MPN iPS cell line undergo EndMT during prolonged *in vitro* culture. The use of isogenic MPN iPS cell lines thus provided a valuable tool to validate findings made in the transgenic murine models, and offered an innovative approach to investigate JAK2V617F-associated endothelial dysfunction.

### The endothelial MPL receptor is required for the development of CVD in mice with JAK2V617F-bearing ECs

The thrombopoietin (TPO) and its receptor MPL are key regulators of hematopoietic stem cell survival, proliferation, and differentiation^40^. MPL is also expressed in various vascular ECs, suggesting its potential involvement in cardiovascular function^41-48^. Notably, JAK2 binds to the intracellular domain of MPL and their interaction is known to facilitate MPL receptor trafficking and stability^40^. Based on these findings, we hypothesized that the JAK2V617F mutation modulates endothelial TPO/MPL signaling, thereby leading to altered vascular endothelial function and contributing to the development of CVDs in JAK2V617F-positive MPNs.

To test this hypothesis, we utilized the MPL^fl^ mice, which carry a GFP reporter to monitor the transcriptional activity of the MPL locus^49^. Flow cytometry analysis revealed robust GFP expression in cardiac and pulmonary ECs, indicating abundant MPL expression in the vascular endothelium of these tissues (Figure 5A).

**Figure 5.**
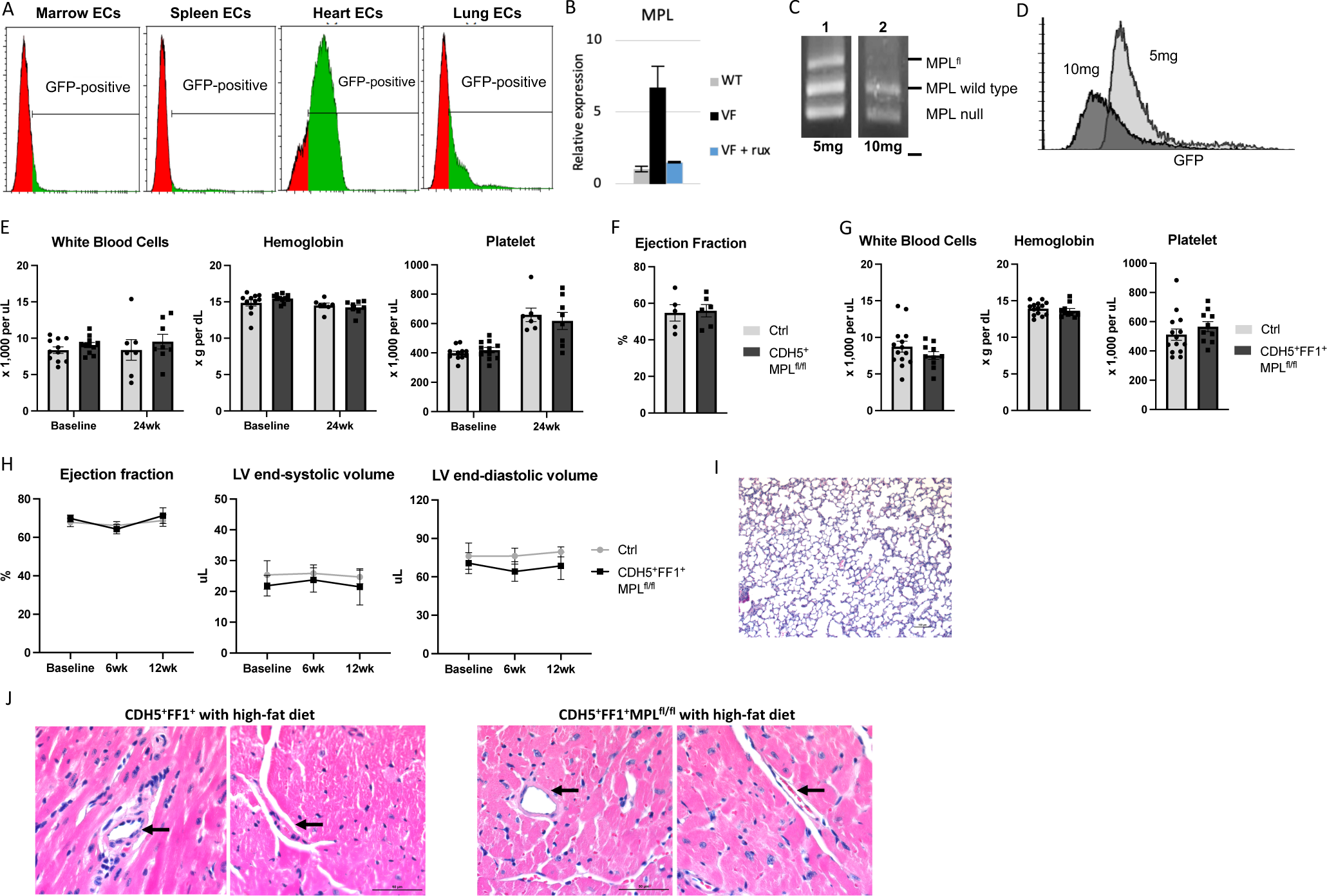
The endothelial MPL receptor is required for JAK2V617F mutant EC-induced cardiovascular dysfunction. (**A**) Representative flow cytometry histogram showing GFP expression in ECs isolated from MPL^fl^ mice, indicating transcriptional activity of the MPL locus. (**B**) MPL expression in wildtype (WT), JAK2V617F mutant (VF), and Ruxolitinib (250nM for 24hrs)-treated JAK2V617F mutant (VF+Rux) lung ECs, measured using real-time quantitative polymerase chain reaction. Gene expression is shown as the fold-change compared to the WT EC expression which was set as ‘1’. (**C**) Genomic DNA PCR analysis of flow cytometry sorted heart ECs (CD45^-^CD31^+^) from CDH5^+^MPL^fl/wt^ mice after 5mg (lane 1) and 10mg (lane 2) tamoxifen induction. 5mg tamoxifen induction resulted in incomplete recombination with residual MPL^fl^ allele, while 10mg tamoxifen resulted in complete recombination with no MPL^fl^ allele left. (**D**) Representative flow cytometry histogram showing EC GFP expression after 5mg and 10mg tamoxifen induction of CDH5^+^MPL^fl/wt^ mice. (**E**) Peripheral blood counts in CDH5^+^MPL^fl/fl^ mice and CDH5^-^MPL^fl/fl^ control mice at baseline and 24 weeks after tamoxifen induction (n=7-11 mice in each group). (**F**) Cardiac ejection fraction in CDH5^+^MPL^fl/fl^ mice and control mice 16-20 weeks after tamoxifen induction (n=5-6 mice in each group). (**G**) Peripheral blood counts in CDH5^+^FF1^+^MPL^fl/f^ and control mice at 16 weeks after tamoxifen induction (n=10-14 mice in each group). (**H**) LV ejection fraction and LV volumes in CDH5^+^FF1^+^MPL^fl/f^ and control mice after high-fat diet challenge (n=5-6 mice in each group). (**I**) H&E staining of lung sections from CDH5^+^FF1^+^MPL^fl/f^ mice after 12-week high-fat diet treatment (magnification 10x). (**J**) Representative H&E staining of intramyocardial coronary arterioles (arrow) in high-fat diet treated CDH5^+^FF1^+^ mice (left) and CDH5^+^FF1^+^MPL^fl/f^ mice (right) (magnification 40x).

Consistent with our previous findings^50,51^, we observed a significant increase in MPL expression in JAK2V617F mutant ECs compared to wild-type ECs. Treatment with the JAK inhibitor Ruxolitinib^52^ effectively abolished this upregulation (Figure 5B), suggesting the involvement of JAK-STAT signaling in JAK2V617F-induced MPL upregulation.

To investigate the functional role of endothelial MPL in normal hematopoietic and cardiovascular functions, we generated VEcad-cre^+/-^MPL^fl^ mice (CDH5^+^MPL^fl^) by crossing VEcadherin-creERT2 mice^29,30^ with MPL^fl^ mice^49^. We evaluated the efficiency of tamoxifen induction in heterozygote CDH5^+/-^MPL^fl/wt^ mice and found that 10mg tamoxifen induction yielded higher recombination efficiency compared to 5mg tamoxifen induction (Figure 5C-D). Therefore, we utilized 10mg tamoxifen to specifically knock down MPL receptor expression in vascular ECs. Following tamoxifen induction, CDH5^+^MPL^fl/fl^ mice maintained normal blood cell counts during a 5-mo of follow-up (Figure 5E). This finding is consistent with a previous report indicating that the endothelial MPL receptor does not significantly contribute to the regulation of steady-state platelet counts^53^. Furthermore, cardiac function was unaffected in CDH5^+^MPL^fl/fl^ compared to control mice (Figure 5F). These results suggest that endothelial MPL is not essential for the normal physiological regulation of blood cell counts and cardiac function.

Previous reports have indicated an increased risk of thrombosis in patients receiving TPO receptor agonist therapy, particularly in the presence of underlying hypercoagulability (e.g., liver disease)^54,55^. These observations suggest that the underlying disease status could influence how TPO/MPL signaling impacts vascular function. To investigate the role of endothelial TPO/MPL signaling in JAK2V617F-induced cardiovascular dysfunction, we generated CDH5^+^FF1^+^MPL^fl/fl^ mice by crossing CDH5^+^FF1^+^ mice with MPL^fl^ mice, allowing for MPL knockdown specifically in JAK2V617F-bearing vascular ECs. In contrast to CDH5^+^FF1^+^ mice, which developed mild thrombocytosis 12 weeks after tamoxifen induction (Figure 1B), CDH5^+^FF1^+^MPL^fl/fl^ mice maintained normal blood counts (Figure 5G) and exhibited preserved cardiac function even after a high-fat diet challenge (Figure 5H). Histological examination did not reveal evidence of pulmonary thrombosis (Figure 5I) or coronary vasculopathy (Figure 5J) in these mice.

In conclusion, our findings suggest that endothelial TPO/MPL signaling plays a significant role in the development of JAK2V617F-induced cardiovascular dysfunction, and targeting this pathway could be a potential therapeutic strategy to prevent or mitigate cardiovascular complications in JAK2V617F-positive MPNs.

### Inhibiting endothelial TPO/MPL signaling suppresses JAK2V617F-induced EndMT in both transgenic murine model and human MPN iPS cell line

Next, we investigated the effects of inhibiting endothelial MPL on the functions of JAK2V617F mutant ECs. As described above, JAK2V617F mutant cardiac ECs from high-fat diet-treated CDH5^+^FF1^+^ mice exhibited reduced angiogenesis (Figure 3B) and decreased expression of EC markers (Figure 3E-F) in comparison to wild-type ECs. In contrast, cardiac ECs isolated from CDH5^+^FF1^+^MPL^fl/fl^ mice (which harbor JAK2V617F-bearing MPL^-/-^ ECs) displayed increased angiogenesis (Figure 6A) and higher levels of VE-cadherin and VEGF-R2 compared to wild-type control cardiac ECs (Figure 6B-C). These findings suggest that inhibiting endothelial MPL can reverse or attenuate the JAK2V617F-induced EndMT process, thereby preventing the development of cardiovascular dysfunction.

**Figure 6.**
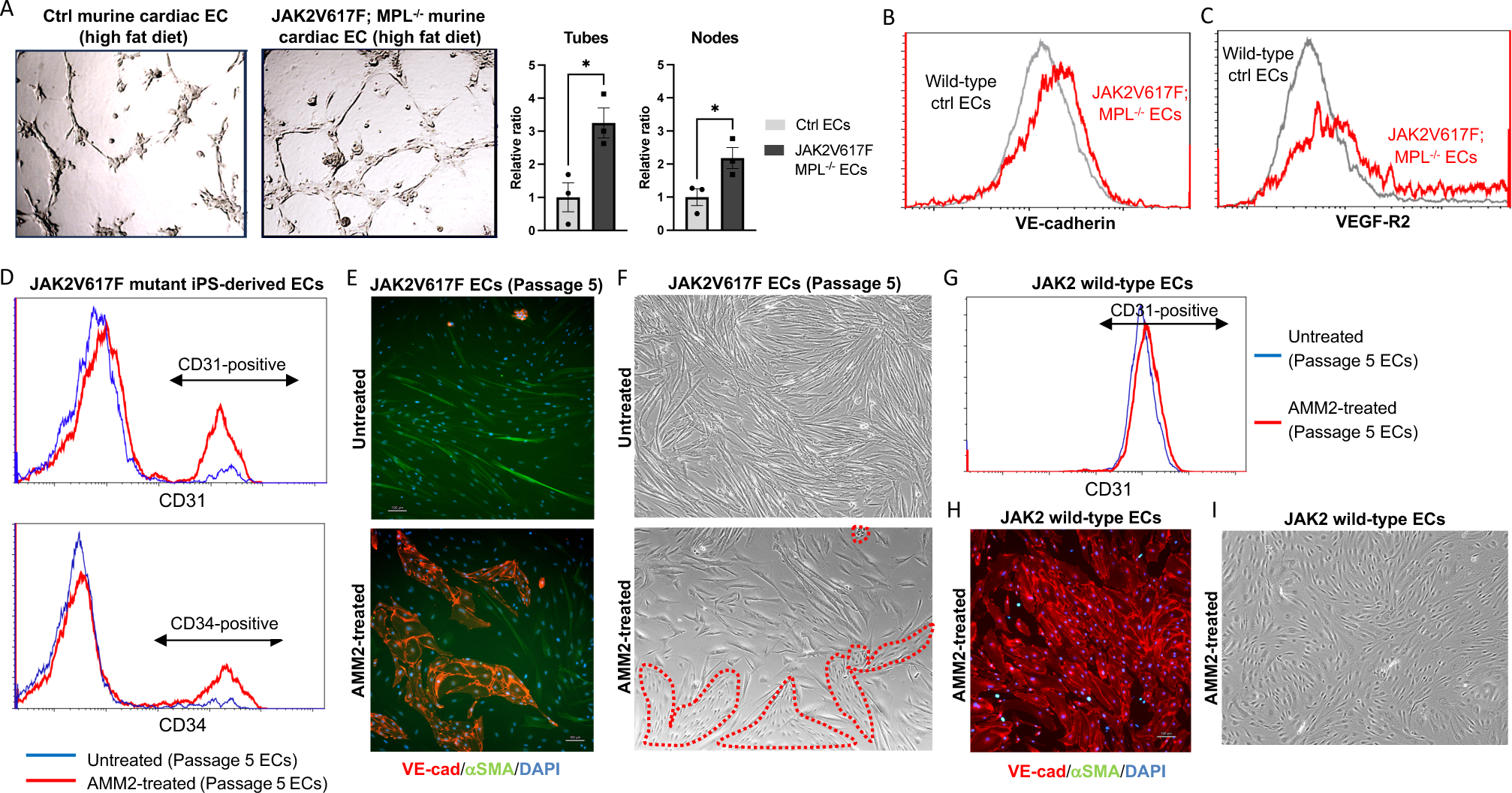
Suppression of JAK2V617F-induced EndMT by inhibition of endothelial TPO/MPL signaling in both a transgenic murine model and a human MPN iPS cell line. (**A**) Tube formation assays demonstrating increased tube formation in JAK2V617F-bearing MPL^-/-^ cardiac ECs (from CDH5^+^FF1^+^MPL^fl/fl^ mice) compared to wild-type cardiac ECs (from CDH5^-^FF1^-^MPL^fl/fl^ control mice) even after high-fat diet treatment. Data are from one of two independent experiments that gave similar results. (**B-C**) Expression levels of VE-cadherin (B) and VEGF-R2 (C) expression in JAK2V617F-bearing MPL^-/-^ cardiac ECs and wild-type cardiac ECs after high-fat diet treatment. Similar results were obtained from two independent experiments. (**D**) Flow cytometric analysis showing increased expression of CD31 and CD34 on AMM2-treated JAK2V617F mutant human ECs 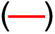 compared to untreated JAK2V617F mutant ECs 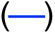. (**E**) Immunofluorescence staining for VE-cadherin (red) and αSMA (green) in JAK2V617F mutant human ECs with (bottom) or without (top) AMM2 treatment. (**F**) Representative bright field images demonstrating that a significant proportion of AMM2-treated passage 5 JAK2V617F mutant ECs retained the cobblestone endothelial morphology (red dotted lines; bottom) compared to untreated mutant ECs, which mostly exhibited a fibroblastoid morphology (top). (**G-I**) AMM2 treatment does not impair the function of PVB1.11-derived JAK2 wild-type ECs, as measured by flow cytometry (G), immunofluorescence staining (H), and microscopic examination (I). For D-I, similar results were obtained from two independent experiments conducted by two different researchers.

To further investigate the protective effect of inhibiting endothelial MPL in JAK2V617F mutant human EC function, we conducted experiments using JAK2 wild-type and JAK2V617F mutant human ECs derived from MPN iPS cell lines (PVB1.11 and PVB1.4). We treated these ECs with the anti-MPL neutralizing antibody AMM2 (100ng/ml) between passage 3 and passage 5 (Figure 3C-E). As shown earlier, untreated JAK2V617F mutant ECs experienced a loss of most EC markers (CD31 and CD34) by passage 5. In contrast, the treatment with AMM2 significantly preserved the expression of these EC markers in JAK2V617F mutant ECs at passage 5 compared to untreated mutant ECs at the same passage (Figure 6D). Immunofluorescence staining revealed that AMM2 treatment increased the number of VE-cadherin-positive cells and reduced the number of αSMA-positive cells in JAK2V617F mutant ECs compared to untreated mutant ECs (Figure 6E). Moreover, microscopic examination demonstrated that while untreated JAK2V617F mutant ECs mostly acquired an elongated fibroblastoid morphology by passage 5, a substantial proportion of AMM2-treated mutant ECs retained the characteristic cobblestone endothelial morphology (Figure 6F). Importantly, AMM2 treatment did not impair the function of JAK2 wild-type human EC, as evidenced by the preserved expression of EC markers and the maintenance of EC morphology on flow cytometry analysis, immunofluorescence staining, and microscopic examination (Figure 6G-I). Taken together, these findings strongly suggest that AMM2 treatment effectively prevented JAK2V617F-induced EndMT in human ECs derived from the MPN iPS cell line.

## Discussions

With the increasing utilization of DNA sequencing, the detection of CHIP is anticipated to increase. As an independent and potent risk factor for CVDs, the assessment and management of individuals with CHIP present a growing challenge to clinicians across various specialties. A key goal for future research is to improve the stratification of CVD risk in individuals with CHIP, enabling more targeted and effective management strategies. Among the various CHIP-associated mutations, the JAK2V617F mutation confers a significantly higher risk of CVDs compared to other mutations^5,7,8^. Vascular ECs play a critical role in the regulation of cardiovascular function and ECs carrying the JAK2V617F mutation have been detected in many patients with MPNs^12,13,16-19^. In this study, we investigated the contribution of JAK2V617F mutant ECs to the development of CVD and explored the underlying mechanisms. Our findings support that JAK2V617F mutant ECs play a significant role in the development of CVD, particularly when exposed to additional stressors such as a high-fat/high-cholesterol diet (Figures 1-2). Both transgenic murine model studies (Figure 3) and experiments utilizing MPN iPS cell lines (Figure 4) consistently demonstrated that the JAK2V617F mutation induces EndMT, a pivotal mechanism implicated in driving endothelial dysfunction and cardiovascular diseases^34-36^. These findings provide valuable insights into the increased risk of CVDs observed in patients with JAK2V617F-positive CHIP and MPNs.

While our study focused on the JAK2V617F mutation, it is important to note that hematopoietic mutations can also be detected in ECs in many hematologic malignancies, such as MPNs, lymphoma, and myeloma^12-19^. To further investigate the presence of these mutations in the vascular endothelium in patients with CVDs, we recently conducted whole exome sequencing on paired blood and vascular samples from four patients with severe coronary artery disease who underwent coronary artery bypass graft surgery. Remarkably, we identified mutations in CHIP-associated genes (JAK2, JAK3, IDH2, U2AF2, ETNK1) in the vascular endothelium isolated from the internal mammary artery or saphenous vein grafts of these patients^56^. These findings are in accordance with previous studies that reported the presence of somatic mutations and clonal expansions in vascular ECs or smooth muscle cells within both healthy and diseased cardiovascular tissues^57-61^. Given the well-established link between CHIP mutations and increased inflammation^62^, which is known to induce EndMT^34-36^, further investigations are needed to understand the involvement of EndMT in other CHIP-associated mutations. These investigations will help elucidate the specific contributions of different CHIP mutations to the diverse risks of CVDs associated with them.

Our study also provided novel insights into the role of endothelial MPL in the development of cardiovascular dysfunction in JAK2V617F-positive MPNs. We demonstrated that inhibiting endothelial TPO/MPL signaling effectively attenuated JAK2V617F-induced EndMT and prevented the development of cardiovascular dysfunction caused by mutant ECs, even in the presence of a high-fat diet challenge (Figures 5-6). Despite extensive research on TPO and its receptor MPL in normal hematopoiesis^63,64^ and their contribution to the neoplastic hematopoiesis of MPNs^65-67^, their roles in cardiovascular complications associated with JAK2V617F-positive MPN remain poorly understood. Our recent work has revealed the critical role of TPO/MPL signaling in the expansion of MPN stem cells within the mutant hematopoietic microenvironment, where major components of the vascular niche (e.g., ECs and megakaryocytes) bear the JAK2V617F mutation^50,51,68^. In conjunction with the findings from our current study, we have uncovered the critical role of endothelial MPL receptor in both the hematopoietic vascular niche (promoting MPN neoplastic stem cell expansion) and the general vasculature (contributing to JAK2V617F-induced cardiovascular dysfunction). Notably, the endothelial MPL receptor is not indispensable for the normal physiological regulation of blood cell counts and cardiac function (Figure 5E), making it a potential therapeutic target for preventing or mitigating cardiovascular complications in JAK2V617F-positive MPNs.

Collectively, this work provides insights for future research and potential clinical application to focus on vascular pathology in the diagnosis, risk stratification, and management of cardiovascular complications associated with hematopoietic mutations.

## Methods

### Experimental mice

JAK2V617F Flip-Flop (FF1) mice, which carry a Cre-inducible human JAK2V617F gene expression driven by the human JAK2 promoter, were provided by Radek Skoda (University Hospital Basal, Switzerland)^27^. Dr. Warren Alexander (Melbourne, Australia) provided the MPL^fl/fl^ mice, which carry a GFP reporter for transcriptional activity of the MPL locus and can undergo a Cre-mediated deletion of the GFP reporter as well as exons 11 and 12, resulting in a null allele (i.e., changing the 388bp MPL^fl^ allele to a 236bp MPL^null^ allele)^49^. Tie2-Cre mice^28^ were obtained from Mark Ginsberg (University of California, San Diego). Cdh5-CreERT2 mice^29^ were purchased from Taconic Biosciences Inc. (Albany, NY, USA). All mice used were crossed onto a C57BL/6 background and housed in a pathogen-free mouse facility at Stony Brook University. No randomization or blinding was used to allocate experimental groups. All animal experiments were conducted in compliance with the guidelines of the Institutional Animal Care and Use Committee.

### Transthoracic echocardiography

Transthoracic echocardiography was performed on mildly anesthetized spontaneously breathing mice (sedated by inhalation of 1% isoflurane, 1 L/min oxygen), using a Vevo 3100 high-resolution imaging system (VisualSonics Inc, Toronto, Canada) as described previously^22^.

### Histology

Hearts and lungs were fixed in cold 4% paraformaldehyde overnight at 4°C while shaking gently. The tissues were then washed multiple times with phosphate buffered saline (PBS) at room temperature to remove paraformaldehyde. Paraffin sections (5-μm thickness) were stained with hematoxylin/eosin (H&E), reticulin, and Masson’s trichrome using reagents and kits from Sigma (Sigma, St. Louis, MO) following standard protocols. Images were taken using a Nikon Eclipse Ts2R Fluorescent Microscope.

### Complete blood counts

Peripheral blood was obtained from the facial vein via submandibular bleeding and collected in EDTA tubes. The samples were then analyzed using a Vetscan Hm5 Hematology Analyzer (Abaxis) to determine the blood count.

### Flow cytometry

All samples were analyzed by flow cytometry using a LSR II flow cytometer (BD Biosciences, San Jose, CA) or a CytoFLEX flow cytometer (Beckman Coulter, Indianapolis, IN, USA). For murine sample flow cytometry analysis: CD45 (clone 104, Cat. 109813, Biolegend, San Diego, CA), CD31 (clone 390, Cat. 102401, BD Biosciences, San Jose, CA), VE-cadherin (clone BV13, Cat. 138006, Biolegend), and VEGF-R2 (clone Avas12, Cat. 136403, Biolegend) antibodies were used. For human sample flow cytometry analysis: CD31 (Clone WM59, Cat. 560984, BD Biosciences), CD34 (Clone 581, Cat. 343515, Biolegend), and CD45 (Clone HI30, Cat. 555485, BD Biosciences) antibodies were used.

### Polymerase Chain Reaction

To detect human JAK2V617F expression, reverse transcription polymerase chain reaction (RT-PCR) was performed using human JAK2-specific primers GAGCAAGCTTTCTCACAAGC and AATTCTGCCCACTTTGGTGC. The amplification of a 530-bp fragment confirms the Cre-mediated expression of the human JAK2 gene in the FF1 transgenic mice.

To verify the recombination status of the MPL locus in MPL^fl/fl^ mice, we conducted PCR using four primers: 5’-CGACCACTACCAGCAGAACA-3’, 5’-GGATGGTGTTGAGGATGGAT-3’, 5’-CCGATAGCTGTGAAGAAGTGG-3’, 5’-ACAGACAACCCCCTGCAGTA-3’. The presence of the MPL^fl^ allele (which carries the GFP reporter) was indicated by a PCR band of 388 bp. Conversely, the MPL^null^ allele resulting from a Cre-mediated deletion of the GFP reporter as well as exons 11 and 12 of the MPL locus was identified by a PCR band of 236 bp.

### Isolation and culture of murine lung and cardiac ECs

Primary murine lung and cardiac EC (CD45^-^CD31^+^) were isolated following a previously described protocol with minor modifications^22,50,69,70^. Briefly, mice were euthanized and the chest was immediately opened through a midline sternotomy. The animal was subjected to cardio-pulmonary perfusion with 30 ml of cold PBS using a 27-gauge needle inserted into the left ventricular apex, with simultaneous drainage through a small incision in the right atrium. The heart and lung tissue were quickly collected and finely minced with scissors. The tissue fragments were then digested in DMEM medium containing 1 mg/mL Collagenase D (Roche, Switzerland), 1 mg/mL Collagenase/Dispase (Roche), and 25 U/mL DNase (Sigma) at 37°C for 2 hours with gentle shaking. The digested tissue suspension was drawn into a 20ml syringe attached to a 16-gauge needle and triturated to obtain a single-cell suspension by passing it through the needle at least 12 times. The homogenate was filtered through a 70μm cell strainer (BD Biosciences, San Jose, CA) and centrifuged at 400g for 5 minutes. The cell pellet was washed with 50ml PBS, and resuspended in an appropriate buffer to first deplete CD45^+^ cells, followed by selection for CD31^+^ cells, using magnetically labeled microbeads according to the manufacturer’s protocol (Miltenyi Biotec, SanDiego, CA). The isolated ECs (CD45^-^CD31^+^) were cultured on 1% gelatin coated plates in complete EC medium, as previously described^69^. No medium change was performed for the first 72 hours to allow EC attachment, followed by medium change every 2-3 days. Cells were re-depleted for CD45^+^ cells when they reached >70-80% confluence, usually after ∼7 days of culture.

### Maintenance and Expansion of Human iPS cells

A pair of human induced pluripotent stem (iPS) cell lines (JAK2 wild-type PVB1.11 and JAK2V617F mutant PVB1.4) derived from the same MPN patient were obtained from Drs. Zhaohui Ye and Linzhao Cheng (Johns Hopkins University)^71^. The iPS cells were cultured on vitronectin-coated dishes (0.5 ug/cm^2^, A14700, ThermoFisher) in Essential 8 Medium (A1517001, ThermoFisher). To improve cell viability, RevitaCell Supplement (A2644501, ThermoFisher) was added in the medium for the first 24 hours post-thaw. When the cells reach 70-80% confluence, they were passaged as small clumps using 0.5mM EDTA.

TRA-1-60 live staining and immunostaining for undifferentiated markers such as NANOG and SSEA4 were performed to confirm the pluripotency of the iPS cells. The karyotyping assay, conducted at the Johns Hopkins Cytogenetics Lab, was used to confirm the cytogenetic stability of the iPS cells. The JAK2V617F mutation status of PVB1.11 and PVB1.4 was verified using a nested allele-specific PCR assay, as described previously^72^. The presence of a 279-bp product indicated the JAK2V617F-positive allele, whereas a 229-bp product indicated the wild-type allele. All cell lines were routinely tested for mycoplasma contamination and were negative throughout this study.

### Endothelial Differentiation of iPS cells

Both JAK2 wild-type (PVB1.11) and JAK2V617F mutant (PVB1.4) iPS cell lines were differentiated into ECs using a modified iPS differentiation protocol adapted from Patsch et al.^38^ and Guadall et al.^39^ (see Figure 4A below). *Day0*: iPS cells were dissociated using Accutase (A6964, Sigma) and seeded on Growth Factor Reduced Basement Membrane Matrix-coated plates (Cat. 356230, Corning) at a density of 30,000 cells/cm^2^ in mTeSR medium with 10uM RevitaCell. *Day1*: the medium was replaced by N2B27 medium, which is consisted of a 1:1 mixture of DMEM:F12 (Cat. 11320082, ThermoFisher) and Neurobasal medium (Cat. 21103049, ThermoFisher) supplemented with N2 (Cat. 17502048, ThermoFisher) and B27 without vitamin A (Cat. 12587010, ThermoFisher), containing 7μM of CHIR99021 (Cat. 4423/10, R&D Systems) and 25ng/mL recombinant human BMP4 (Cat. 314-BP, R&D Systems), at ∼0.7ml of medium per cm^2^ of cultured area. *Day 4*: the medium was replaced with StemPro-34 SFM (Cat. 10639011, ThermoFisher) containing 200ng/mL Vascular Gndothelial Growth Factor (VEGF)-165 (Cat. 100-20, Peprotech) and 2μM Forskolin (Cat. ab120058, Abcam), at ∼0.3ml of medium per cm^2^ of cultured area. *Day6-7*: cells were dissociated using Accutase and separated using human CD144 Microbeads (Cat. 130-097-857, Miltenyi). Isolated CD144^+^ cells were plated at a density of ∼15,000-25,000 cells/cm^2^ on human Fibronectin-coated dishes (Cat. 354008, Corning) in StemPro-34 SFM supplemented with 50ng/mL VEGF-165. *Day8-9*: the medium was replaced with StemPro-34 SFM supplemented with 50ng/mL VEGF-165 and 8μM SB431542 (Cat. 1614/10, R&D systems). From *Day12* onwards, ECs were cultured in EGM-2 medium (Cat CC-3162, Lonza) supplemented with 8μM SB431542. The medium was replaced every other day. Cells at confluence were dissociated using 0.05% Trypsin-EDTA (Cat. 25300054, ThermoFisher) and cryopreserved in freezing medium consisting 90% FBS and 10% DMSO for future experimental use. The iPS-derived wild-type ECs maintained their EC identity up to at least passage 8, although cell growth was significantly slowed after passage 6.

### Assays to examine endothelial cell in vitro angiogenesis

The EC tube formation assay was performed to evaluate angiogenesis *in vitro*^22,50,51,73^. Matrigel^®^ matrix (Cat. 354234, Corning) was thawed overnight at 4°C and kept on ice until use. A pre-chilled 96-well culture plate was prepared, and 50μl of Matrigel was added to each well. The plate was then incubated at 37°C for 30 minutes to allow gelation. Next, 1 × 10^4^ ECs in 100ul of corresponding EC medium were added on top of the Matrigel. The tube formation process was observed at different time points, including 2, 4, 6, 8, and 24 hours, and images captured using a phase-contrast microscope at both 4x and 10x magnifications (AMEX-1200, ThermoFisher). Quantification of the tube formation was performed using the ImageJ^®^ Angiogenesis Analyzer (National Institute of Health, Bethesda, MD) by counting the number of tubes and nodes in 4-5 non-overlapping areas.

### Immunophenotyping of human iPS-derived ECs

The cells were washed with PBS and fixed with 4% paraformaldehyde for 10minutes at room temperature with gentle rocking. After removing the paraformaldehyde, the cells were washed three times with PBS (5 minutes each). Next, the cells were permeabilized with 0.5% Triton X-100 in PBS for 10minutes at room temperature, followed by three washes with PBS. Cells were then blocked with 5% BSA at room temperature for 30-60min prior to antibody staining. The blocking medium was used for all subsequent antibody staining steps. The cells were incubated overnight at 4ºC with mouse anti-human αSMA (Cat. 14-9760-82, Invitrogen: 1:500 dilution) and goat anti-human CD144 (Cat. AF938, R&D Systems; dilution 1:250). After three washes with PBS, the cells were incubated with donkey anti-goat IgG conjugated to AF555 (Cat. A21432, ThermoFisher; 1:500 dilution) for 30minutes to 1 hour at room temperature. Following another round of washing (three times), the cells were incubated with goat anti-mouse IgG conjugated to AF488 (Cat. A11001, ThermoFisher; 1:500 dilution) for 30minutes to 1 hour at room temperature. Cells were then washed and incubated in DAPI (Cat. D9542, Sigma; 1:5000 dilution of 10mg/ml stock) for 10 minutes at room temperature. Finally, the cells were washed three times for 5 minutes each with PBS and imaged using a Nikon Eclipse Ts2R Fluorescent Microscope. If necessary, samples can be stored in PBS, protected from light, at 4ºC for at least 1 week for re-imaging purposes.

### Ruxolitinib and AMM2 treatment

JAK2 wild-type and JAK2V617F mutant primary murine lung ECs (passages 2-4) were cultured in complete EC medium^22,50,69,70^ and treated with either DMSO or 250nM Ruxolitinib (Selleckchem®) for 24 hours. After treatment, cells were harvested and total RNA was isolated using RNeasy Mini kit (Qiagen) for gene expression analysis.

Human iPS-derived ECs were cultured in EGM-2 medium supplemented with 8μM SB431542 as described above. To inhibit TPO/MPL signaling, cells were treated with 100ng/ml of anti-MPL antibody (AMM2, IBL-America). The expression of EC markers was monitored after each passage using both flow cytometry analysis and immunofluorescence staining.

### Quantitative real-time quantitative PCR Analysis

The TaqMan® Gene Expression Assay of MPL (Mm00440306_g1, ThermoFisher) was used for real-time quantitative polymerase chain reaction to verify differential expression of MPL on an Applied Biosystems 7300 Real Time PCR System (ThermoFisher). Values obtained were normalized to the endogenous Actin beta (Mm00607939_s1, ThermoFisher) gene expression and relative fold changes compared to control samples was calculated by the 2^ΔΔCT^ method. All assays were performed in triplicate.

### Transcriptomic analysis of cardiac ECs using RNA sequencing

RNA sequencing of JAK2 wild-type and JAK2V617F mutant cardiac ECs (CD45^−^CD31^+^) (n = 3 mice in each group) were described previously^22^. Gene set enrichment analysis (GSEA) was performed using the complete list of genes according to their differential expression.

### Statistical analysis

Data analysis and graph creation were performed using Excel software (Microsoft) and GraphPad Prism (GraphPad Software, La Jolla, CA). Statistical analysis was conducted using two-tailed unpaired Student’s *t* test unless otherwise specified. A P-value of <.05 was considered statistically significant. The results are presented as mean ± SEM (standard error of the mean).

## Acknowledgements

This research was supported by the VA Merit Award BX003947 and BX005584 (H.Z.) and NIH R01 HL134970 and R01 CA266294 (H.Z.).

## Author Contributions

H. Zhang conducted transgenic murine model studies and contributed to experiments using iPS-derived ECs; N.K. and K.M. conducted experiments involving iPS-derived ECs; S.L. contributed to iPS cell culture protocol and prepared the figures; H. Zheng assisted with the analysis of cardiovascular histology; H. Zhan conceived the projects, designed and performed the experiments, acquired and analyzed the data, interpreted the results, and wrote the manuscript.

